# Comprehensive network modeling approaches unravel dynamic enhancer-promoter interactions across neural differentiation

**DOI:** 10.1101/2024.05.22.595375

**Authors:** William DeGroat, Fumitaka Inoue, Tal Ashuach, Nir Yosef, Nadav Ahituv, Anat Kreimer

**Affiliations:** Center for Advanced Biotechnology and Medicine, Rutgers, The State University of New Jersey, 679 Hoes Lane West, Piscataway, NJ 08854, UAS; Institute for the Advanced Study of Human Biology (WPI-ASHBi), Kyoto University, Kyoto, Japan; Department of Electrical Engineering and Computer Sciences and Center for Computational Biology, University of California, Berkeley, 387 Soda Hall, Berkeley, CA 94720, USA; Department of Systems Immunology, Weizmann Institute of Science, 234 Herzl Street, Rehovot 7610001, Israel; Chan-Zuckerberg Biohub, 499 Illinois St, San Francisco, CA 94158, USA; Department of Systems Immunology, Ragon Institute of MGH, MIT, and Harvard Institute of Science, 400 Technology Square, Cambridge, MA 02139, USA; Department of Bioengineering and Therapeutic Sciences, University of California, San Francisco, 513 Parnassus Ave, CA 94143, USA; Institute for Human Genetics, University of California, San Francisco, 513 Parnassus Ave, CA 94143, USA; Department of Biochemistry and Molecular Biology, Rutgers, The State University of New Jersey, 604 Allison Road, Piscataway, NJ 08854, USA

**Keywords:** Functional Genomics, Gene Regulation, Neural Development, Computational Genomics

## Abstract

**Background:** Increasing evidence suggests that a substantial proportion of disease-associated mutations occur in enhancers, regions of non-coding DNA essential to gene regulation. Understanding the structures and mechanisms of regulatory programs this variation affects can shed light on the apparatuses of human diseases.

**Results:** We collected epigenetic and gene expression datasets from seven early time points during neural differentiation. Focusing on this model system, we constructed networks of enhancer-promoter interactions, each at an individual stage of neural induction. These networks served as the base for a rich series of analyses, through which we demonstrated their temporal dynamics and enrichment for various disease-associated variants. We applied the Girvan-Newman clustering algorithm to these networks to reveal biologically relevant substructures of regulation. Additionally, we demonstrated methods to validate predicted enhancer-promoter interactions using transcription factor overexpression and massively parallel reporter assays.

**Conclusions:** Our findings suggest a generalizable framework for exploring gene regulatory programs and their dynamics across developmental processes. This includes a comprehensive approach to studying the effects of disease-associated variation on transcriptional networks. The techniques applied to our networks have been published alongside our findings as a computational tool, E-P-INAnalyzer. Our procedure can be utilized across different cellular contexts and disorders.

## Background

Enhancers are cis-regulatory elements (CREs) essential to gene regulation. Ranging between ten to 10,000 base pairs (bps) in length [1], these CREs contain motifs, clustered recognition sites for transcription factors (TFs) [2]. Enhancers have proven arduous to locate despite their abundance in the non-coding genome. Enhancers interact with promoters to mediate their effects on transcription, often by regulating non-proximal genes from variable distances [3]. This contrasts with promoters, CREs that initiate transcription from known locations [4]. Enhancers are highly cell-type-specific regulatory elements, which adds another barrier to their identification and validation [5]. There is an emerging consensus that mutations associated with human disease in genome-wide association studies (GWAS) are concentrated in these enhancer regions [6], highlighting their importance to understanding gene regulation and applications in medicine.

Previously, we collected high-throughput assay for transposase-accessible chromatin (ATAC-seq), chromatin immunoprecipitation (ChIP-seq) for H3K27, massively parallel reporter assay (MPRA), and RNA-seq datasets from seven time points during the differentiation of human embryonic stem cells (hESCs) into neural progenitor cells (NPCs) [7]. This was within a 72-hour window, with samples collected at hours 0, 3, 6, 12, 24, 48, and 72. Using these datasets, we assembled seven time point-specific enhancer-promoter interaction networks (E-P-INs). ATAC-seq, indicating open chromatin regions, ChIP-seq for H3K27ac, acting as a marker for active enhancers, and RNA-seq, representing the transcriptome, combined with generalized Hi-C datasets, serve as the essential components for these networks. Input into the Activity-By-Contact (ABC) model, these datasets integrate to build networks of predicted enhancers and target promoters [8]. After constructing these networks, information about the enhancers within them was extracted using a series of novel methods: differential expression revealed time point specificity, Girvan–Newman clustering demonstrated the substructures through which enhancers regulate genes, variant analyses revealed enrichment for neurodevelopmental and neuropsychiatric disorders, and TF overexpression and MPRA datasets were used to validate our networks.

Overall, our analysis of these E-P-INs provides a framework for exploring gene regulatory programs and their dynamics across developmental processes, including a comprehensive approach for mapping disease-associated genetic variants to transcriptional networks.

## Results

### Reconstructing enhancer-promoter interactions with multi-omics datasets

Examining shifts in the intricate mechanisms of regulatory programs across seven stages of neural differentiation necessitated designing a systematic procedure to generate E-P-INs. Our E-P-INs are highly modular bipartite graphs populated with communities of enhancers and promoters; edges represent interactions between these elements. These networks were designed for cross-condition comparability, referencing the same atlas of enhancers and promoters (see “Methods”). Our E-P-INs capture the temporal programs of gene regulation within our seven time points exclusively, omitting ubiquitous CREs that are unchanging.

Eight E-P-INs were created using the ABC model: seven time-point-specific networks and a single collapsed E-P-IN containing all edges found across these time points. Here, we report statistics from the collapsed network to summarize the structures of the seven time-point-specific E-P-INs. Our E-P-INs contained 3,133 unique promoters and 1,628 enhancers. This suggests most interactions in these E-P-INs, consisting of differentially active and temporal CREs, are non-redundant, with an average of 1.92 genes regulated through a single enhancer. The mean length of enhancers in our E-P-INs was 1,047.83 bps; this aligns with previous literature [1, 9]. Examining sample composition, we found consistent interactions within our time-point-specific networks. The smallest E-P-IN (48 hours) contained 2,134 interactions, totaling 13.02% of our collapsed network’s edges. The largest E-P-IN (24 hours) included 15.68% of edges in the collapsed network, amounting to 2,569 unique enhancer-promoter links. Tab. ST1 in “Supplementary Material” contains our time-point-specific and collapsed E-P-INs.

### Enrichment in neural differentiation signals

Initial investigations of our E-P-INs showed characteristics relevant to early neural differentiation through enrichment and temporal analyses. Our E-P-INs demonstrated enrichment for biological processes pertinent to the cellular context and temporal changes from the start of neural induction to the 72-hour time point.

Gene Set Enrichment Analysis (GSEA) was performed utilizing the 500 most frequently occurring promoters from each E-P-IN [10]. The Genomic Regions Enrichment Annotations Tool (GREAT) was used to examine enrichment in our enhancer regions [11, 12]. See “Methods” for an in-depth explanation of GSEA and GREAT’s methodologies. We report the results from our collapsed E-P-IN for brevity; time-point-specific results are available in Tab. ST2 and Tab. ST3 in “Supplementary Materials”. The most significant results from GSEA and GREAT are reported in Fig. 1A and Fig. 1B, respectively. “GOMF_TRANSCRIPTIONAL_REGULATOR_ACTIVITY,” “primary neural tube formation,” and “ventral spinal cord interneuron specification” are examples relevant to neural differentiation. We note that GSEA results relate to transcription due to the number of promoters being tested and their recurrence across the networks. GREAT considers all enhancers in our E-P-INs. A concentrated GSEA testing set contributes to the difference in statistical magnitudes between GSEA and GREAT. These findings support our E-P-INs’ biological relevance.

**Figure 1.**
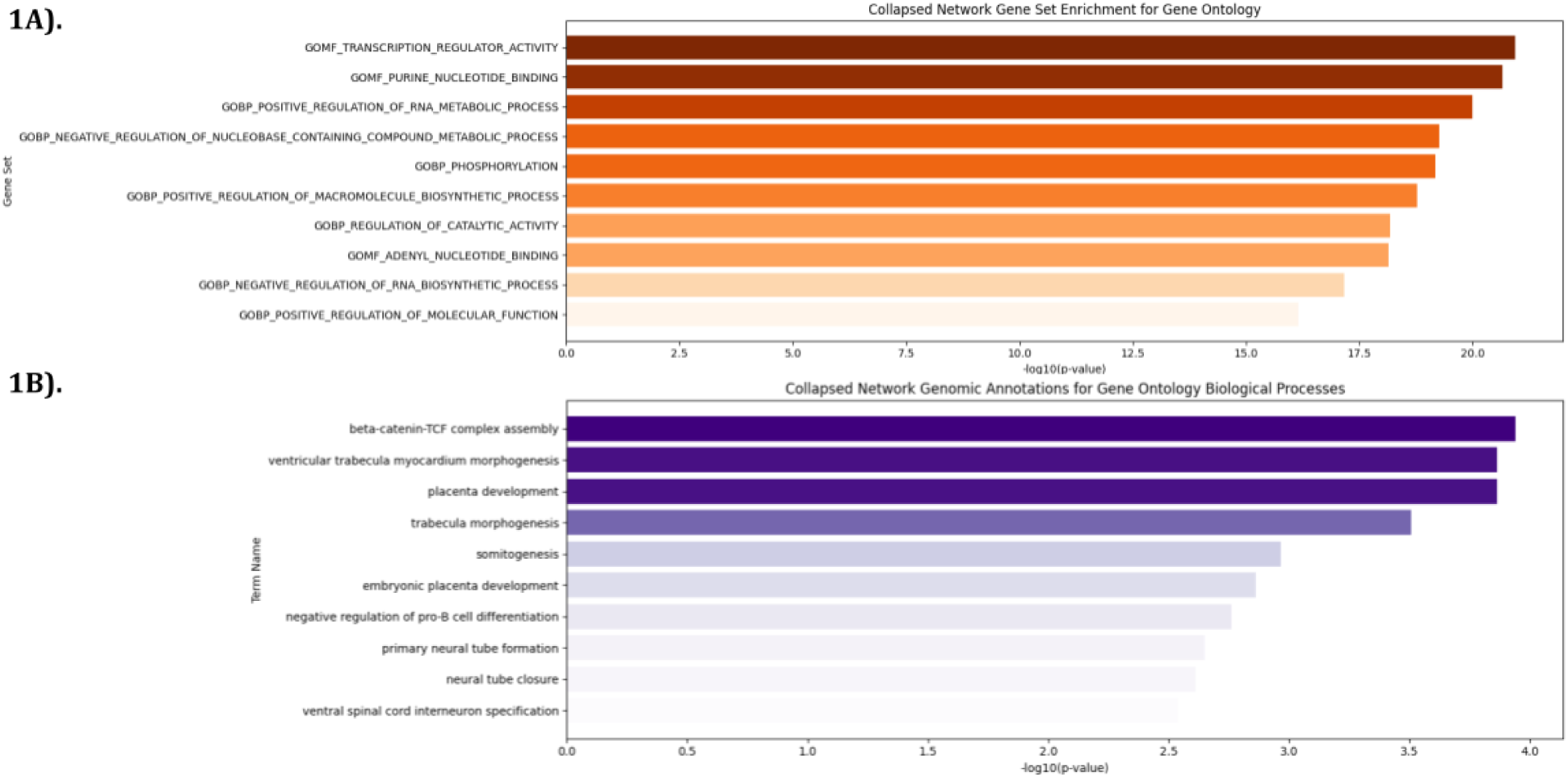
Enrichment analyses demonstrate the E-P-INs’ abilities to capture neural differentiation. 1A) GSEA outcomes from the collapsed network’s 500 most occurring promoters reveal biological processes, cellular components, and molecular functions essential to transcriptional regulation. 1B) GREAT outcomes detailing enhancer enrichment show biological processes related to cell differentiation and neural development.

Next, Jaccard similarities between the CREs (i.e., enhancers and promoters) and enhancer-promoter interactions (E-P-Is) were computed to compare our seven time-point-specific E-P-INs. Fig. 2A, 2B, and 2C present Jaccard similarities in enhancers, promoters, and E-P-Is, respectively. Here, we show that these networks become more dissimilar with time. Additionally, this analysis revealed that time points before 48 hours show high similarity compared to our 48-hour and 72-hour E-P-INs; these latter networks also show close resemblance. These results indicate our E-P-INs exhibit temporal trends of neural differentiation.

**Figure 2.**
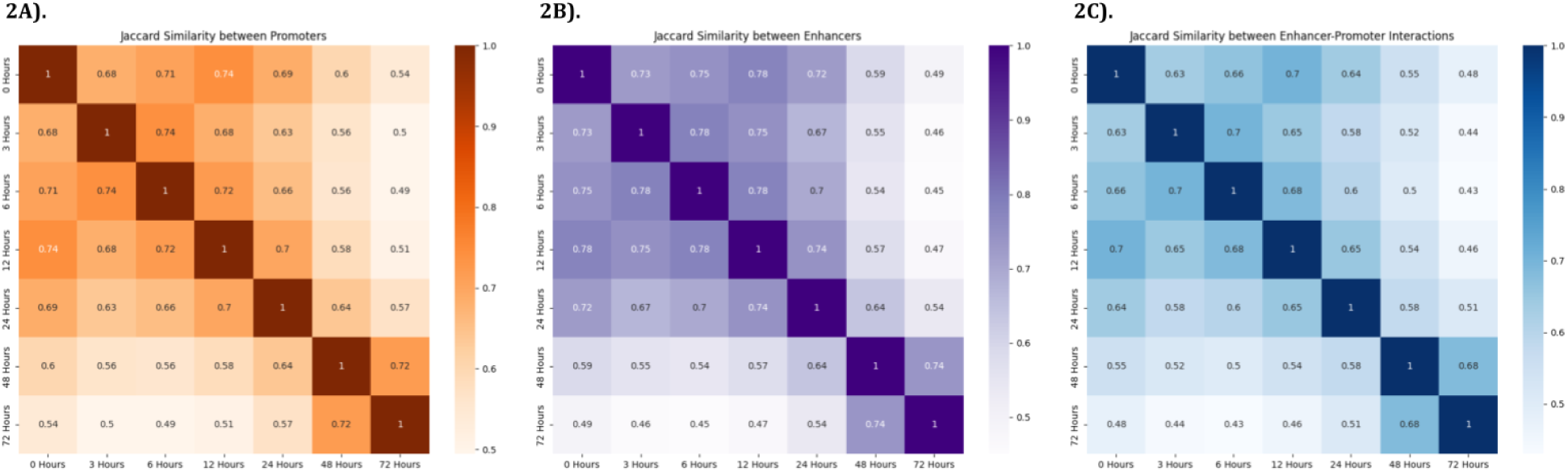
Temporal analysis revealed the E-P-INs exhibit changes related to neural differentiation. 2A) Jaccard similarities detail the overlap between sets of promoters across our seven time-point-specific networks, with clustered formation in the earliest five time points. 2B) Jaccard similarities detailing differences in time-point-specific sets of enhancers reveal temporal trends similar to those described for promoters. 2C) Jaccard similarities demonstrating changes in the E-P-Is across neural differentiation. Instances where connections between enhancers and promoters change are more temporal than either the present/absence of enhancers or promoters individually.

### Clustering reveals regulatory substructures

We identified four distinct substructures defining the modes through which enhancers regulate genes. Here, Girvan-Newman clustering was utilized to partition our bipartite graph into distinct communities [13]. The highly modular nature of our E-P-INs informed this approach, as enhancers and promoters tied together through gene regulation formed dense clusters in our networks. Fig. 3 provides a graphical representation of our collapsed E-P-IN to illustrate the formations of these substructures. Substructures fall into two distinct classifications: redundant (R) and non-redundant (NR). The four substructures are a single enhancer regulating a single gene (1NR; Fig. 3A), multiple enhancers regulating a single gene (1R, Fig. 3B), a single enhancer regulating multiple genes (2NR; Fig. 3C), and multiple enhancers regulating multiple genes (2R; Fig. 3D). Fig. SF1 in “Supplementary Materials” contains a full view of our collapsed E-P-IN [14].

**Figure 3.**
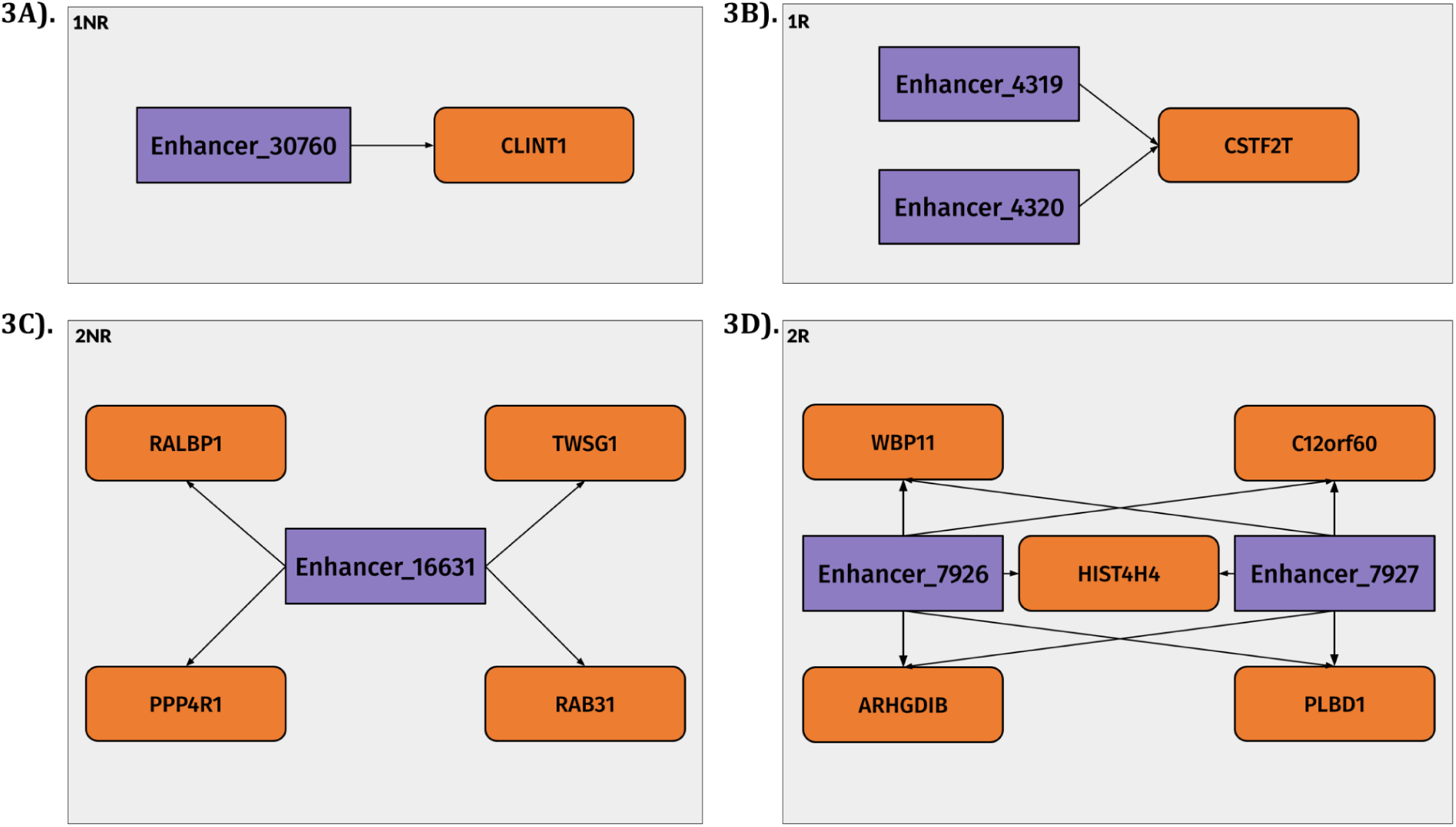
Graphical representations demonstrate the composition of gene regulatory substructures. 3A) offers an example of a 1NR substructure. 3B) shows a community of E-P-Is characterized as 1R. 3C) demonstrates the 2NR substructure. 3D) depicts a cluster labeled as 2R.

We implemented this clustering approach for each of the seven time-point-specific E-P-INs and the collapsed E-P-IN. Fig. 4A shows the four distinct substructures as a legend, and Fig. 4B displays the substructure distribution in the seven time-point-specific E-P-IN, calculated by dividing the total number of communities by the number of communities allocated to each specific substructure. This distribution remains consistent, with slight fluctuations, across our seven time points. The 1NR and 2NR substructures constitute the majority of our time point-specific networks, averaging 81.9% composition across the seven E-P-INs. This skew towards NR substructures stems from the dynamic and temporal filtering used to reconstruct the networks.

**Figure 4.**
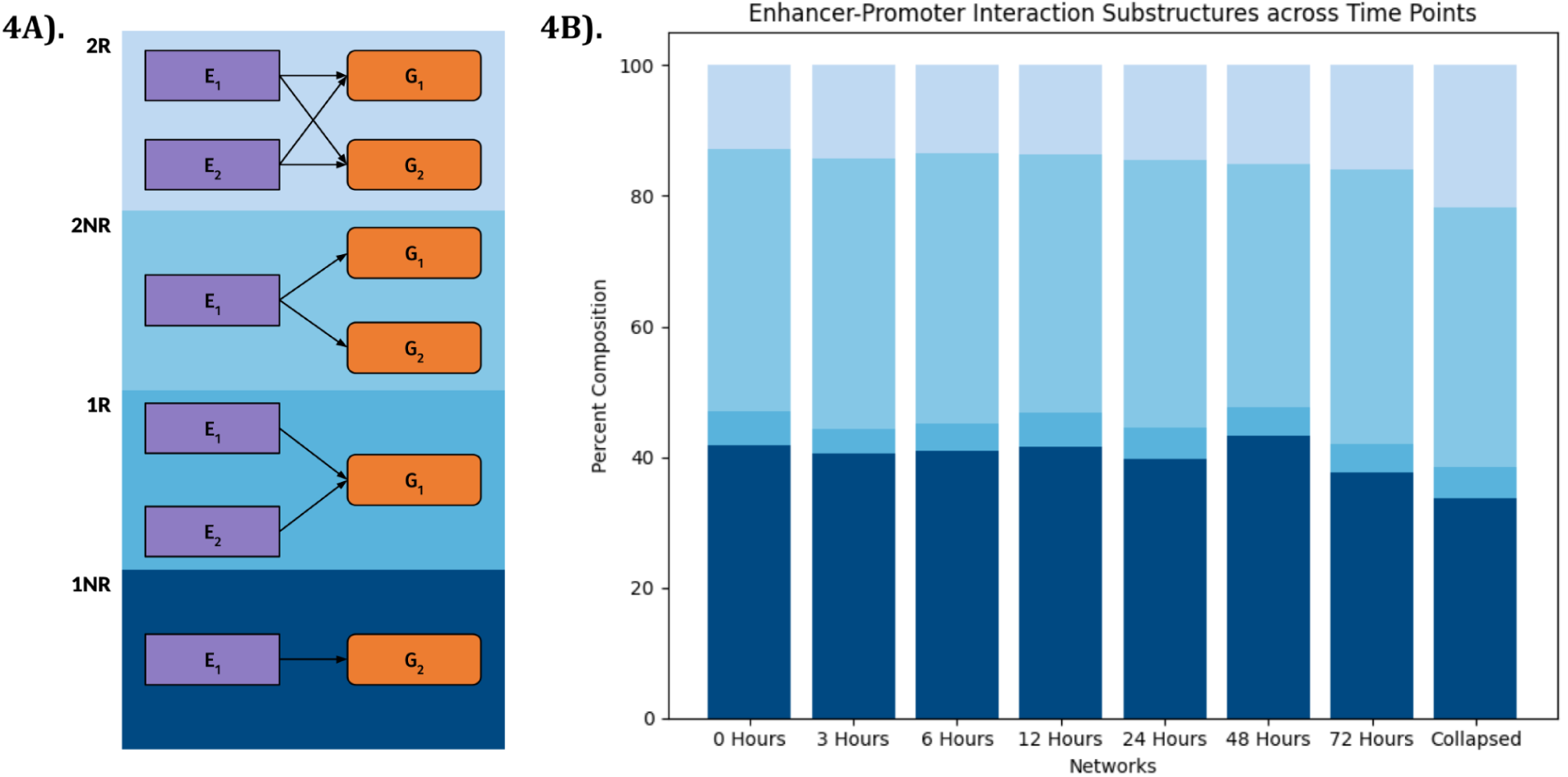
Substructures across the seven time-point-specific E-P-INs demonstrate minimal temporality. 4A) is a legend that links substructures to their appropriate section on the chart. 4B) shows the distribution of substructures across the seven time-point-specific and collapsed E-P-INs. Redundant substructures comprise a smaller portion of these networks than those non-redundant.

We calculated average counts of enhancers, promoters, and E-P-Is in these regulatory substructures using our collapsed E-P-IN. In 1R communities, an average of 2.16 enhancers regulate a single gene; in 2NR communities, 3.29 genes, on average, are regulated through a single enhancer. Interestingly, the 2R substructure is denser, averaging 2.55 enhancers, 4.57 genes, and 7.72 E-P-Is per community. Fig. SF2 compiles histograms showing the distribution of these values. Our investigations could not reveal significant links between gene expression and a promoter’s number of edges with enhancers.

### Enrichment for disease-associated variants

We hypothesized that our time-point-specific E-P-INs would exhibit enrichments associated with neurodevelopmental and neuropsychiatric disorders [15]. Utilizing datasets containing 250,000 de novo autism spectrum disorder (ASD) single nucleotide variants (SNVs) from both patients (cases) and healthy siblings (controls) [16], 817 ASD-associated genes, 252 developmental delay-associated genes (DD) [17], and 34,914 GWAS-sourced SNVs from neurological/neuropsychiatric disorders (i.e., schizophrenia (SCZ), bipolar disorder (BD), major depressive disorder (MDD), Parkinson’s disease (PD), and Alzheimer’s disease (AD)), we implemented a permutation test to quantify an empirical p-value signifying enrichment; our null distribution in these permutation tests was the ABC model’s unfiltered outputs. Eight tests were employed on each E-P-IN: ASD-associated SNVs, ASD-associated genes, DD-associated genes, SCZ-associated SNVs [18], BD-associated SNVs [18], MDD-associated SNVs [19], PD-associated SNVs [20], and AD-associated SNVs [21].

With the significance threshold set at an empirical p-value = 0.05 and 1,000 permutations ran, five of our eight categories demonstrated significant enrichment in our E-P-INs: ASD-associated genes and SNVs, DD-associated genes, and SCZ-associated and BD-associated SNVs. Tab. ST4 in “Supplementary Material” contains a complete overview of the statistics for disease-associated variant enrichment. MDD-associated SNVs, PD-associated SNVs, and AD-associated SNVs were deemed non-significant. These findings suggest our E-P-INs capture enhancers relevant to neural differentiation; SCZ and BD are disorders with neurodevelopmental origins [22, 23]. Out of the 1,592 enhancers considered in our collapsed E-P-IN, 140 contain either a case or control de novo variant for ASD, with case variants favored; 27 contain a GWAS-sourced SNV for a neurological/neuropsychiatric disorder. Of 3,133 genes, 208 are associated with ASD, and 41 are associated with DD. ASD was the most significantly enriched among diseases/disorders, not exceeding a p-value < 0.001 in examinations of associated SNVs and genes. Fig. 5A depicts the results of the ASD-associated de novo SNV permutation test, while Fig. 5B presents the ASD-associated gene permutation test results. Further confirming the enrichment of ASD, we devised an additional permutation test, utilizing the dynamic and temporal E-P-Is as the null distribution. We found that ASD-associated genes demonstrate significant interaction with enhancers containing an ASD-associated SNV, computed at an empirical p-value < 0.001. ASD enrichment arises from a common set of genes and SNVs across our seven time-point-specific E-P-INs, indicating temporal patterns of regulation. Fig. SF3 in “Supplementary Material” shows Jaccard similarities between sets of ASD-associated CREs in our networks.

**Figure 5.**
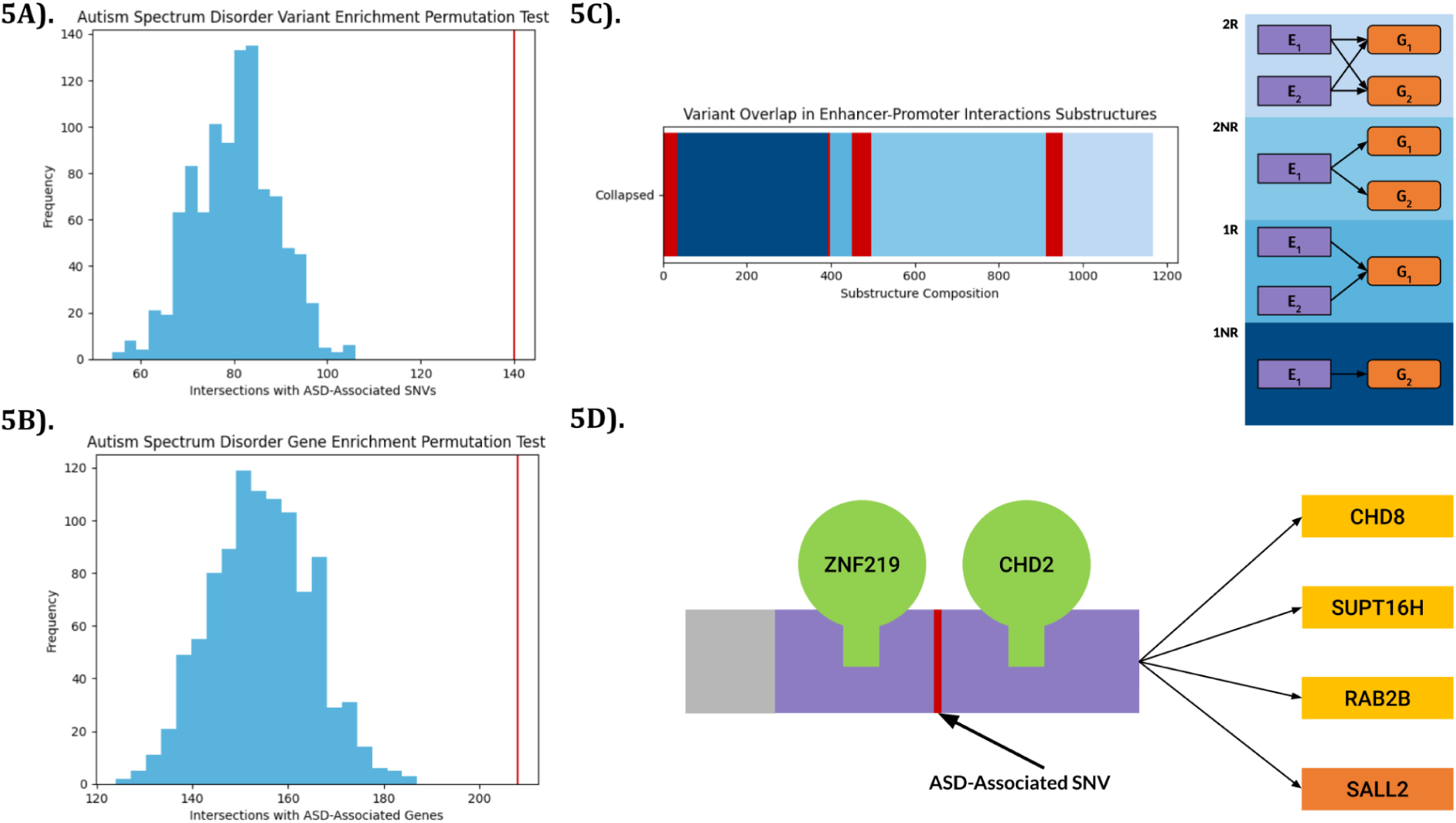
ASD enrichment across E-P-INs is substantial. 5A) demonstrates significant enrichment for ASD-associated de novo SNVs in the collapsed network. 5B) details the enrichment of ASD-associated genes, showing permutation test outcomes. 5C) displays the number of communities within each substructure containing ASD-associated variation. The communities with variants are shaded red at the start of each substructure’s bar. 5D) provides an example of a predicted enhancer region, binding ASD-associated TFs CHD2 and ZNF219, that regulated CHD8, SUPT16H, and RAB2B. Clinical reports provide cases where a mutation in chromosomal region 14q11.2, the location of this enhancer, involved these genes and caused ASD.

We mapped our ASD-associated de novo SNVs to the communities created with the Girvan-Newman clustering algorithm [13]. The distribution of SNVs was consistent with that of substructures in our E-P-INs; 8.42% of 1NR communities contained variation, 10.34% of 1R communities, 10.15% of the 2NR communities, and 15.75% of 2R communities. Fig. 5C details the number of communities in each substructure that contain ASD-associated variants. Prior research has suggested enhancer redundancy as a counteracting mechanism to variation that would disrupt gene transcription [24]. Examining the deleteriousness of the SNVs found in R substructures compared to NR substructures could provide further proof of this claim. Fig. 5D gives an example of the susceptibility of non-redundant substructures, illustrating a case where an ASD-associated SNV fell within a predicted enhancer. Both TFs, ZNF219 [25] and CHD2 [26], have been previously found to be linked to ASD. The enhancer regulates three genes (i.e., CHD8, SUPT16H, and RAB2B); these genes have been linked to ASD through differential expression, clinical variant analyses, and gene knockouts, respectively [27, 28, 29]. Mutations in chromosomal region 14q11.2, where the predicted enhancer is located, have been shown to involve these genes and cause ASD [30, 31].

### Validating enhancers with transcription factor overexpression

Integration of TF data into our seven time-point-specific networks enabled us to begin verification of E-P-Is and gain a more in-depth understanding of these enhancers’ role in gene regulation [32]. First, we used Find Individual Motif Occurrences (FIMO) [33] with motifs from ENCODE [34] and JASPAR [35] databases to find TFs predicted to bind to the sequences of our enhancers. Our p-value threshold for FIMO was 0.00001. Previous studies suggest that tens of TFs can bind to an enhancer [1]; this was in line with our findings. Enhancers in our E-P-INs average 12.30 unique TFs bound to them. Examining our original network, produced with the ABC model, prior to the robust filtration process reported in “Reconstructing Enhancer-Promoter Interactions with Multi-omics Datasets,” we found a decrease in the number of unique TFs per enhancer (7.33). In total, 1183 unique TFs were predicted in our E-P-INs using FIMO. Next, we examined which TFs were more commonly predicted to bind to our enhancers; returning to our ABC network prior to filtration, CTCF, a TF essential to E-P-Is, was bound to the largest number of enhancers. Differing in our dynamic E-P-INs, SP9, a TF necessary for neuronal development [36], was the most common. SP9 binds to 1,222 of the 1,628 enhancers in our time-point-specific networks. These findings suggest that our framework captures TF time point specificity alongside E-P-Is.

Previously, we overexpressed TFs (i.e., BARHL1, IRX3, LHX5, OTX1, OTX2, and PAX6) [7] followed by RNA-seq in later-stage hESC-derived NPCs to detect their target genes. Using DESeq2 for differential expression analysis [37], we determined which genes were significantly affected following a TF’s overexpression. Using these results, we investigated whether enhancers predicted to bind these TFs regulated the differentially expressed genes (DEGs) downstream; downregulated DEGs were not considered in this analysis (see “Methods). Using Fisher’s exact test, p-values were assigned to each TF-binding enhancer, quantifying their relation to the overexpressed TF’s regulon. We found enhancers predicted to bind each of the six overexpressed TFs where a substantial proportion of downstream promoters regulate DEGs. Here, the p-values of the most significant enhancers are reported per TF: BARHL1 = 6.3e-11, IRX3 = 1e-6, LHX5 = 1.6e-4, OTX1 = 3.1e-4, OTX2 = 5.1e-4, PAX6 = 4.2e-9. In addition to this analysis, we designed a permutation test to determine if the proportion of downstream DEGs in enhancers predicted to bind overexpressed TFs was different than in enhancers that are not. Upregulated and downregulated DEGs were examined together. We found that all TFs scored an empirical p-value < 0.001.

PAX6 overexpression is sufficient to induce hESCs into neural lineage [38]. We report an additional enhancer predicted to bind PAX6 here. This enhancer’s targets in our E-P-IN are enriched with DEGs downregulated during PAX6 overexpression. Only PHKG2, one of nine targets for this enhancer, was upregulated during PAX6 overexpression. In the PAX6-binding enhancer found to have a significant proportion of upregulated DEGs downstream, DDA1 was the sole downregulated target DEG. This suggests a pattern of enhancers regulating directional gene sets and a possible method of examining TF antagonism [39]. Fig. 6A details the permutation test outcomes from PAX6. Fig. 6B illustrates the most significant TF-binding enhancer predicted to bind PAX6 when examining upregulated DEGs. Fig. 6C illustrates the most significant TF-binding enhancer predicted to bind PAX6 when examining downregulated DEGs.

**Figure 6.**
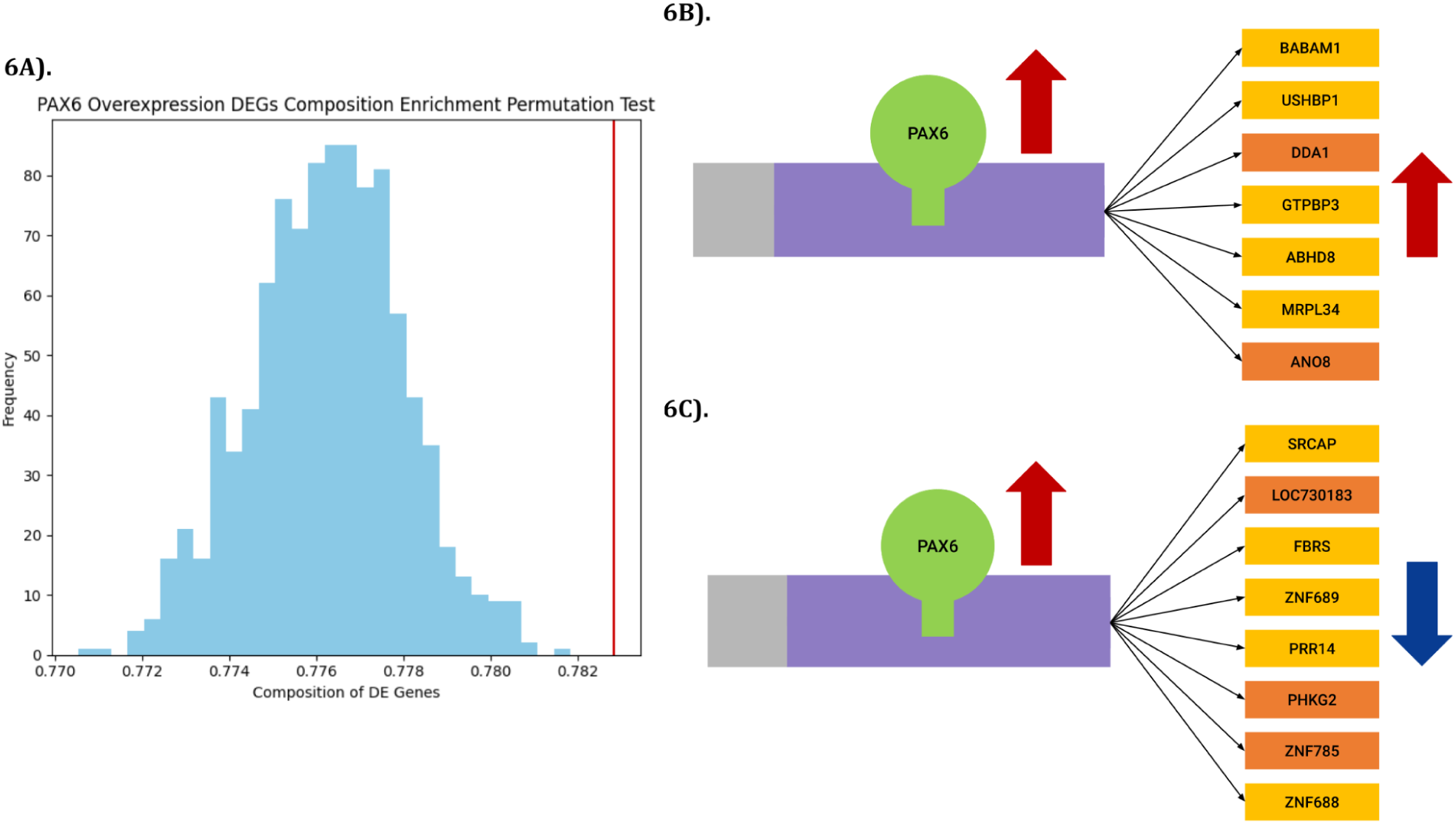
Enhancers predicted to bind PAX6 more commonly regulated DEGs when PAX6 is overexpressed. 6A) shows outcomes from permutation testing supporting the claim that enhancers binding PAX6 regulate DEGs more often than enhancers without PAX6 motifs. 6B) details an enhancer region predicted to bind PAX6 that regulates upregulated DEGs. Genes shown in lighter colors are DEGs when PAX6 is overexpressed. 6C) details an enhancer region predicted to bind PAX6 that regulates downregulated DEGs.

### Validating enhancers with massively parallel reporter assays

In MPRAs, a synthetic construct, each comprising a candidate regulatory DNA sequence, a minimal promoter, and a unique transcribable DNA barcode, is introduced into cells; these candidates are assumed to be able to regulate the transcription of these barcodes. Following this insert, RNA and DNA sequencing are performed to evaluate each barcode’s RNA-to-DNA ratio [40]. MPRAs allow us to measure the activity of thousands of regulatory sequences at once. We used our previously collected perturbation MPRA datasets [7, 41, 42] to validate our predicted enhancers. Perturbation MPRA is a strategy for demonstrating the regulatory effects of TF-binding motifs. Here, nucleotides in a motif are randomly scrambled. We then analyze the change in regulatory activity of the DNA sequence being tested to see the effects of TF binding. We examined E-P-IN enhancers for strict MPRA overlap, creating a subsetted network. This consolidated E-P-IN allowed for an in-depth investigation into TFs, as we computed a novel metric for TF x TF correlation using enhancer binding, genes regulated downstream, and time-point-specific function.

Our subsetted MPRA-based E-P-IN had strict inclusion criteria for edges. An MPRA-tested sequence had to intersect each included enhancer. Additionally, these sequences also needed to perturb a TF that matched a FIMO-predicted motif for that enhancer (See “Methods”). This E-P-IN contained 45 enhancers and 82 promoters, a ratio comparable to the collapsed E-P-IN (1.82 to 1.92). Applicable edges matching the E-P-INs’ criteria were found more often at some time points. A total of 195 edges in the 6-hour network matched, while only 69 in the 72-hour E-P-IN were found.

We devised a permutation test to investigate whether MPRA-validated enhancers were significantly enriched in our temporal, dynamic networks (see “Methods”). Using the ABC model networks as the null distribution, we computed an empirical p-value < 0.001 from 1,000 permutations, showing that proven enhancers are more common in networks of E-P-Is after our filtration methods are utilized.

We devised a TF x TF correlation metric, incorporating enhancers that these TFs are predicted to bind, genes they regulate downstream, and the time points at which the TFs are functional. The metric is based on Jaccard similarities; scores between 0.0 and 1.0 are created for the three components that were previously mentioned. Tab. ST5 contains matrices for these component-specific scores and our TF x TF metric. Fig. 7 shows a heatmap of similarity scores subjected to hierarchical clustering to reveal groups of correlated TFs. TF families, such as the NKX-homeodomain TFs [43] and IRF TFs [44], cluster together, supporting our approach’s ability to reveal TFs working in tandem. Analyses were performed on our MPRA-subset networks to provide interpretable results, detailing a group of enhancers binding TFs validated using experimental methods.

**Figure 7.**
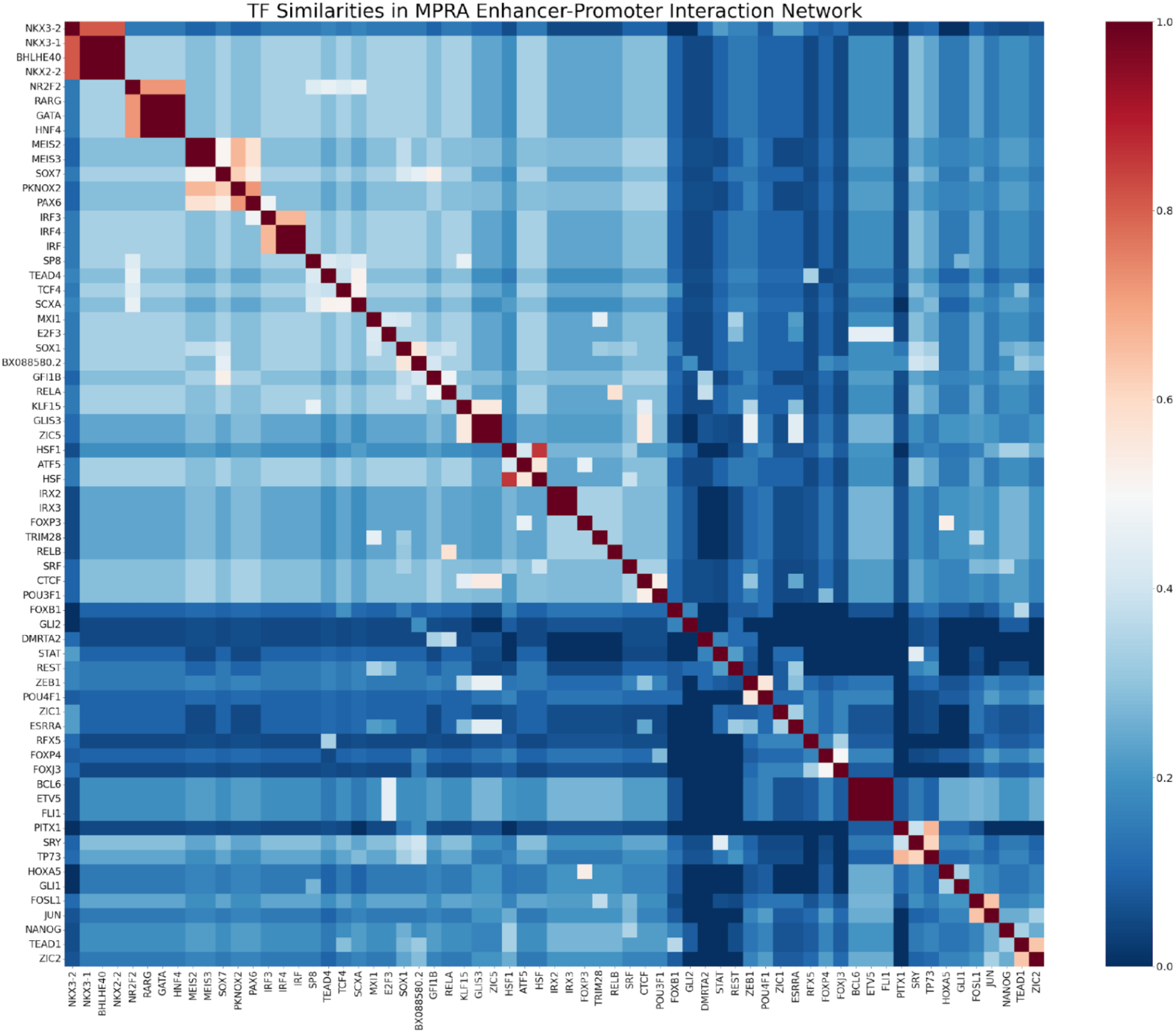
TF x TF matrix, stylized as a heatmap and organized using hierarchical clustering, displays our Jaccard-based similarity metric for transcription factors in our subsetted E-PN derived from MPRA datasets.

## Discussion

Understanding gene expression and the architecture of E-P-Is has the potential to transform human health. Three distinct challenges confront researchers attempting to fill knowledge gaps regarding enhancers: locating enhancers, finding their target genes, and contextualizing their functions [45]. We present a comprehensive computational approach to navigate these questions.

The ABC model has proven to be a well-performing method for predicting E-P-Is, with standard use across the field. Additional layering with perturbation MPRAs allows for built-in motif and variant validation [41, 46]. Unraveling the TFs that bind enhancers is pivotal to their identification, as TF binding is linked to enhancer activity [1]. Developing models for E-P-I predictions using MPRA data to define candidate enhancers and score their activity could tremendously increase the strength of the E-P-I predictions, allowing for better capture of cell-type-specific elements. Our E-P-INs subset with MPRA data showed TF per enhancer counts, average bp lengths, and gene-to-enhancer counts in line with previous scientific findings. MPRA, paired with TF overexpression data and differential expression methods, offers a robust framework for the validation of E-P-Is, helping us understand concrete links between genes and these non-coding regions.

Contextualizing enhancer function is tied to temporal functions, shifting connections between enhancers and genes and enhancers that remain dormant during specific stages in development. Using E-P-INs, as we proposed, allows comparison of CREs between different stages of development to be done with ease. Variant information can also be mapped directly into these E-P-INs and their substructures, allowing us to mold disease networks from model systems. This framework is generalizable, and with sufficient datasets, our analysis can be repeated across biological systems. E-P-INAnalyzer has been developed alongside our analysis as a computational tool that allows other scientific researchers to model biological networks, perform clustering to reveal substructures, integrate TFs and variants, and search for network enrichment. This tool has been made available on our GitHub, found in “Availability of data and materials.” With the same ATAC-Seq and ChIP-Seq for H3K27ac needed to run the ABC Model and RNA-seq, E-P-INAnalyzer performs these analyses, returning condition-specific networks. TF and variant datasets are optional for building networks.

## Conclusion

In this study, we have provided a robust, generalizable framework for comparing E-P-INs across multiple conditions and cellular contexts. We have developed techniques to verify computationally predicted enhancers using TF overexpression and MPRA. We have also demonstrated the use of these E-P-INs in understanding the mechanisms by which disease-associated variation perturbs regulatory networks. E-P-INAnalyzer, our computational tool, allows researchers to answer similar questions about their model system of interest; this tool creates E-P-INs and provides methods to understand them.

## Methods

### Reconstructing enhancer-promoter interactions with multi-omics datasets

Our E-P-INs’ architecture was composed using the Python [47] and R [48] programming languages. These languages enabled us to create comparable networks.

Assembling an E-P-IN is contingent on the ABC model [8], which uses epigenetic datasets to define candidate enhancers and predict their interactions with promoters. The ABC model relies on these straightforward biological notions: enhancers are distinguished through signals denoting potential for transcriptional regulation (ATAC-seq, ChIP-seq), and frequent promoter contact (Hi-C) indicates an edge. Here, we ran the ABC model using the authors’ recommended parameters. Peak-calling was performed using MACS2 on ATAC-seq [49]; these peaks were resized to 500 bps around their center. The 150,000 peaks containing the most read counts were maintained for downstream analysis. Hi-C data averaged from 25 cell types was utilized at a 5-Kb resolution (ENCFF134PUN). We generated ABC networks for each of the seven time points we sequenced.

We constructed time-point-specific E-P-INs. First, a consolidated atlas was generated, correcting for sequencing disparities across samples. Enhancers located less than 100 bps away from each other were concatenated, leaving their time-point-specific programs of regulation intact. These enhancers were then assigned searchable reference labels. Enhancers in this atlas were filtered according to two non-exclusive criteria. Regions that showed significant differential expression (p-values = 0.05) and temporal expression (p-value = 0.01) in our ATAC-seq and ChIP-seq H3K27ac were included in our E-P-INs. DESeq2 conducted differential expression [37] across possible pairings of time points, and ImpulseDE2 was used to determine temporal expression [50]. Next, we included regions in the 95th percentile of counts. Promoters were processed similarly using our RNA-seq datasets. DEGs (p-value = 0.05) that demonstrated temporal expression (p-value = 0.01) were included in our E-P-INs. Promoters not included using the earlier mentioned threshold in the highest 5% of TPM values were included. This was done to capture highly expressed cellular processes during neural differentiation; 283 of the 19,825 candidate genes for our reconstructed E-P-INs are placed using the TPM threshold alone. Note that we utilized the ABC model with a blacklist of genes that are ubiquitously expressed across all cell types. Edges in our E-P-INs must contain both a selected enhancer and promoter. Using this technique, we compiled our seven time-point-specific E-P-INs and the collapsed network. The package pandas was utilized to store and manipulate these networks [51].

### Enrichment in neural differentiation signals

We characterized our E-P-INs using two distinct forms of investigation. Using GSEA and GREAT, we performed enrichment analyses of enhancers and promoters, respectively [10, 11, 12]. We captured their biological processes, cellular components, and molecular functions. Using Jaccard similarities, we observed our E-P-INs morph over time.

GSEA is a functional-class-scoring method for pathway analysis. In GSEA, a pre-defined gene list is used to evaluate whether gene sets linked through shared biological functions are overrepresented in one of two compared conditions. It focuses on the collective patterns of these sets rather than changes in single genes. We used GSEA to examine the Gene Ontology (GO) collections; we provided the computational tool with the 500 most occurring promoters within our networks. GREAT is an overrepresentation-analysis method. GREAT provided GO enrichment for the nearest genes to our enhancer regions. GREAT compares the proportions of gene sets in these enhancers to those in a background model, in this case, the entire human genome, to evaluate enrichment. In both cases, we report p-values from these tests. Visualizations are created using the Python packages Matplotlib [52] and Seaborn [53]. These visualizations use the -log(p-value) of GSEA and GREAT outcomes, positing the most significant GO in the chart’s longest and darkest bars.

Jaccard similarities are statistics used to measure overlaps between two sets. Jaccard similarities are a percentage of the intersection in two sets compared to the union. These scores serve as the foundation for our cross-network comparisons. All-inclusive sets of promoters, enhancers, and their edges from our E-P-INs are used to calculate Jaccard similarities, comparing the time-point-specific E-P-INs across these three metrics.

### Clustering reveals regulatory substructures

Revealing substructures tied to an enhancer’s regulation of genes necessitated using an algorithm for clustering built around betweenness centrality. This metric divides graphs using a path of their shortest connections, separating dense clustering in communities [13]. This approach, combined with our bipartite graphs’ high modularity, creates subnetworks between enhancers and promoters that we believe mimic regulatory profiles. Girvan-Newman clustering was employed using Python’s iGraph implementation [54].

After creating communities, we categorized them into four distinct categories: 1NR, 1R, 2NR, and 2R substructures. These categories represent redundant and non-redundant regulatory programs. 1NR entails a single gene regulated through a sole enhancer. 1R substructures see a single gene regulated by multiple enhancers. 2NR substructures have one enhancer responsible for regulating multiple genes. 2R communities consist of promoters and enhancers, all with redundancies, interacting in a large network. Distributions reported are percentages of substructures over the total communities. We reported average enhancer, promoter, and E-P-Is per community in each substructure.

### Enrichment for disease-associated variants

Enrichment for disease-associated variation was assessed using in-house permutation testing and distinct sets of SNVs and genes. These permutation tests allowed for randomizable attempts and performed 1,000 iterations. SNV tests are employed using the BedTools Suite [55].

ASD-associated genes and DD-associated genes [17] both had an empirical p-value < 0.001 in their respective permutation tests. These tests determine whether our E-P-INs, built using dynamic and temporal CREs, demonstrate enrichment for these disease-associated genes. Using the ABC Model promoters before differential and temporal filtration as our null distribution, we extract unique gene lists equal in size to those of the E-P-IN we are testing in each iteration. The empirical p-value increases as higher counts of disease-associated genes are found in randomized null lists than in the list sourced from our E-P-IN. An analogous test is conducted for enhancers, as randomized lists of enhancers from our ABC model networks serve as a null distribution, intersecting with variants. Each iteration that counts more variant hits than our target E-P-IN increased the empirical p-value.

Additionally, we examined ASD-associated SNV enrichment through two additional methods. First, we calculated the percentage of communities in each substructure that contained an AS-associated de novo mutation. Next, we devised a permutation test to evaluate if ASD-associated SNV-containing enhancers interact with ASD-associated promoters with significance. We shuffled edges between enhancers and promoters in our network using filtered E-P-Is as the null distribution. We calculated an empirical p-value, which increased when more E-P-Is in our null distribution contained an enhancer with an ASD-associated SNV regulating an ASD-associated promoter. The permutation test employs 1,000 permutations.

### Validating enhancers with transcription factor overexpression

TF binding was predicted in our enhancers using FIMO [33], a computational tool in ENCODE’s MEME suite. We sourced position weight matrices from ENCODE [34] and JASPAR databases [35]. The consolidated atlas of enhancers was formatted as a BED file, converted to FASTA using the hg19 reference genome [56], and input into FIMO. Using a significance threshold of 1e-5, we located TF binding sites (TFBSs) predicted to fall in our enhancers. TFs were then extrapolated from TFBSs.

We used Fisher’s exact test to measure the distribution of DEGs and non-DEGs in each enhancer that bound one of the six TFs for which we collected overexpression RNA-seq datasets. Here, DEGs found downregulated (log2 fold change < 0) were not considered [7, 57, 58, 59, 60, 61, 62]; these TFs function primarily as transcriptional activators. The p-value generated was used to find the enhancers’ relation to the overexpressed TF. DEGs were found at a p-value below 0.01 in our DESeq2 analysis.

We developed a permutation test to examine if enhancers predicted to bind overexpressed TFs regulated DEGs more frequently than enhancers that did not. We calculated the promotion of DEGs in both TF-binding and non-binding enhancers. Next, the DEG label is shuffled across genes, and the proportions are recalculated. The empirical p-value denotes the number of occurrences where the shuffled DEGs are found downstream of predicted non-TF-binding enhancers more often than enhancers thought to bind the overexpressed TFs. The permutation test iterates 1,000 times before generating an empirical p-value. Here, the directionality of differential expression was not utilized; our analysis is focused on examining how these TFs drive neural differentiation. Examining genes impacted by co-binding or antagonistic TFs contributes to a better understanding of TFs’ role in our model system.

### Validating enhancers with massively parallel reporter assays

We recently published perturbation MPRA datasets [7, 41, 42], systematically perturbing binding motifs across hundreds of enhancers and validating their functionality at the same time points in early neural differentiation as our E-P-INs’ model. We used this data to validate TFs predicted to bind by FIMO and enhancers in our networks [7]. This validation comes with stringent criteria, through which we validated a small subset of our E-P-INs.

Our MPRA-filtered E-P-INs overlap with perturbation-based datasets. MPRA-tested regions need to intersect with candidate enhancers included in assembled E-P-INs; this intersection needs to be in the proper context; the enhancer must have regulatory activity simultaneously when the MPRA is perturbing a motif. Finally, the perturbed motif also needs to be supported using our FIMO analysis.

We created a permutation test to investigate whether MPRA-validated enhancers were enriched in our filtered networks. We calculated the number of enhancers overlapping MPRA-tested sequences in 1000 random instances of our ABC model networks; we calculated an empirical p-value to show the count of these permutations that surpassed the number of overlaps in our collapsed E-P-IN.

Using this smaller network, we derived a similarity metric that is Jaccard-based. We generated sets of enhancers, downstream genes, and time points that TFs bind to, regulate, and are functional. These sets create individual Jaccard scores between our TFs, which are averaged at equal weights into a composite score. These scores are then hierarchically clustered with SciPy [63].

## Discussion

### Ethics approval and consent to participate

Not applicable.

### Consent for publication

Not applicable.

### Availability of data and materials

Computation methods necessary to reproduce this study, including the tool E-P-INAnalyzer, can be found at <https://github.com/KreimerLab/E-P-INAnalyzer>.

### Funding

This work was supported in part by the National Institute of Mental Health grant R00MH117393-05 (A.K.).

### Authors contributions

A.K. conceived and designed the analysis, collected data, wrote the manuscript, and developed analysis ideas and tools. W.D. performed analysis, wrote the manuscript, and developed analysis ideas and tools. T.I., T.A., N.Y., and N.A. contributed analysis ideas. All authors read and approved the final manuscript.

## Acknowledgments

Not applicable.

## Abbreviations

CRE: Cis-regulatory element
bps: base pairs
TFs: transcription factors
GWAS: genome-wide association studies
ATAC-Seq: assay for transposases-accessible chromatin
ChIP-Seq: chromatin immunoprecipitation
MPRA: massively parallel reporter assay
hECSs: human embryonic stem cells
NPCs: neural progenitor cells
E-P-INs: enhancer-promoter interaction networks
ABC: activity-by-contact
GSEA: gene set enrichment analysis
GREAT: genomic regions enrichment annotations tool
E-P-I: enhancer-promoter interaction
R: redundant
NR: non-redundant
ASD: autism spectrum disorders
SNV: single nucleotide variant
DD: developmental delay
SCZ: schizophrenia
BD: bipolar disorder
MDD: major depressive disorder
PD: Parkinson’s disease
AD: Alzheimer’s disease
FIMO: find individual motif occurrences
DEG: differentially expressed gene
GO: gene ontology
BSs: bindings sites

