## Supplementary Materials for "Comprehensive network modeling approaches unravel dynamic enhancer-promoter interactions across neural differentiation"

### Supplementary Material

Figure SF1: Bird's-eye view of collapsed E-P-IN reveals highly modular, bipartite nature.

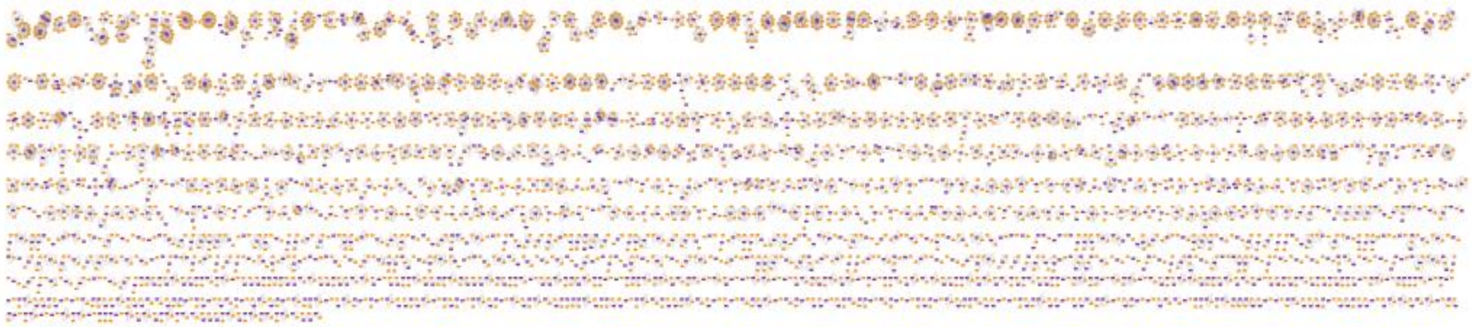

Figure SF2: Distributions of enhancers, genes, and E-P-Is across relevant substructures.

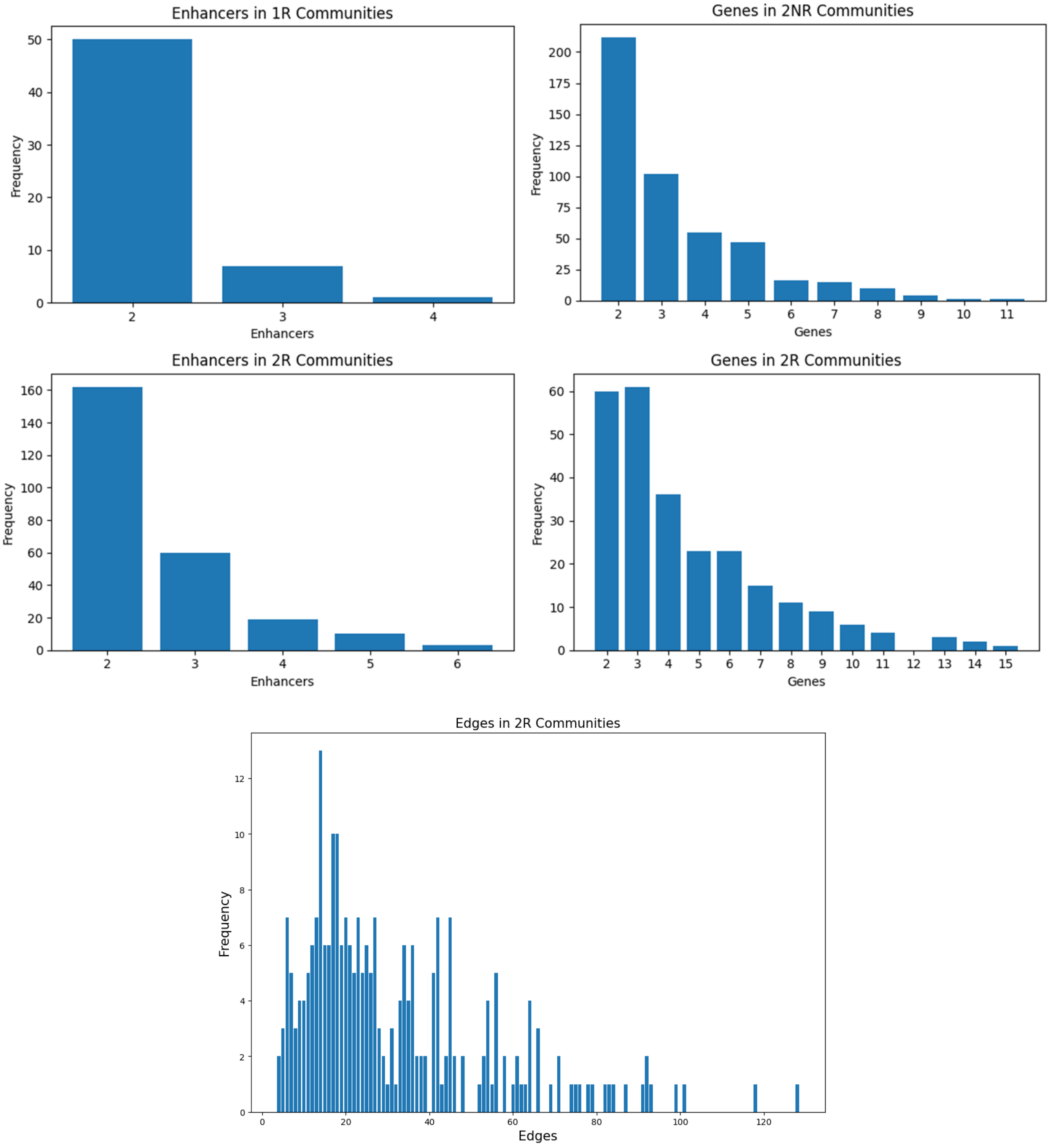

Figure SF3: Jaccard similarities between ASD-associated gene sets in time-point-specific E-P-INs.

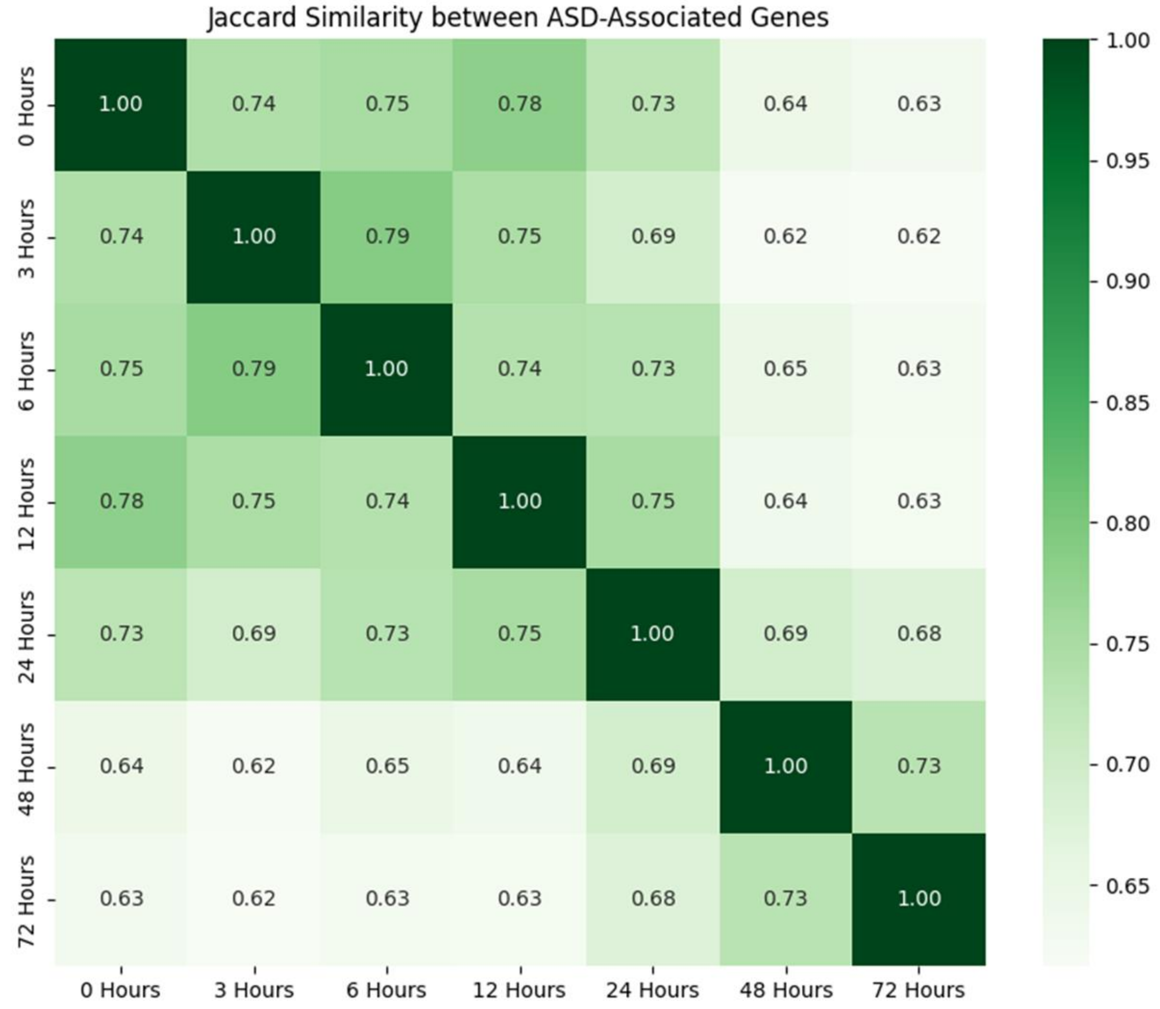

Supplementary Tables:

Table 1: **E-P-INs containing enhancers, promoters, and edges.** Tables containing seven time-point-specific E-P-INs, a collapsed E-P-IN, and an MPRA-validated E-P-IN.

File: Genome-Biology_EPIs_ST1.xlsx

Table 2: **GSEA findings in E-P-INs.**

File: Genome-Biology_EPIs_ST2.xlsx

Table 3: **GREAT findings in E-P-INs.**

File: Genome-Biology_EPIs_ST3.xlsx

Table 4: **Disease-associated variant enrichment in E-P-INs.** Empirical p-values from our permutation tests for time-point-specific and collapsed E-P-INs.

File: Genome-Biology_EPIs_ST4.xlsx

Table 5: **TF similarities.** Three submatrices detailing enhancer, gene, and time point similarities between TF pairs in an MPRA-validated E-P-IN and the composite TF by TF similarity score matrix.

File: Genome-Biology_EPIs_ST5.xlsx
